# In vivo calcium imaging identifies functionally and molecularly distinct subsets of tongue-innervating mechanosensory neurons

**DOI:** 10.1101/2022.02.11.480171

**Authors:** Yalda Moayedi, Shan Xu, Sophie K Obayashi, Benjamin U Hoffman, Gregory J Gerling, Ellen A Lumpkin

## Abstract

Mechanosensory neurons in the mouth provide essential information to guide feeding and speech. How classes of oral mechanoreceptors contribute to oral behaviors is not well understood; in particular, the functional properties of lingual mechanoreceptors remain elusive. Previous work identified putative mechanosensory endings in the tongue with novel morphologies; how these fit into current knowledge of mechanosensory neuron classification is not known. To identify functional classes of lingual mechanosensory neurons, we used *in vivo* calcium imaging of trigeminal ganglia. We first investigated calcium responses of tongue-innervating trigeminal neurons to thermal and mechanical stimulation (e.g., pressure, fluid flow, temperature changes). We found that around 17% of neurons responded to pressure, and that these pressure responders were significantly larger than neurons that only responded to temperature changes. To further investigate the cadre of functionally distinct mechanosensory neurons, we tested responses to brushing and sustained pressures and found that brush-sensitive neurons comprise the majority of tongue-innervating mechanosensory trigeminal neurons. Qualitatively, mechanosensory neurons responded to pressure with distinct kinetics, suggesting the presence of multiple classes of mechanoreceptors. To determine the number of classes, we developed an unbiased multi-layer hierarchical clustering approach to classify calcium response characteristics to pressure stimulation. This approach revealed that mechanosensory neurons displayed five distinct stimulus-response profiles to pressure. Classes include neuronal populations with sustained, transient, high-threshold, and negative responses to force as well as neurons that responded only to brushing. Analysis of cluster representation in transgenic animals with only subsets of labeled neurons reveals molecular markers of clusters and end organ structures. These studies are amongst the first to determine the functional properties of low-threshold mechanosensory neurons innervating the mouse tongue.

## Introduction

Trigeminal sensory neurons innervating the oral cavity provide sensory feedback during a myriad of everyday behaviors including feeding, self and social grooming, and vocalization. For example, during feeding, somatosensory neurons encode the thermal, chemical and textural features of foodstuff, which signal freshness and caloric content (Canon et al., 2021; Klein, 2019; Lemon, 2017; Schobel et al., 2014; Yu et al., 2012). To chew and swallow without injury, animals rely on mechanosensory neurons embedded in oral and upper airway mucosa (Inoue et al., 1989; Moore et al., 2014; Thexton et al., 1980; Travers & Norgren, 1986). Mechanosensory neurons in the oral cavity are equally important for social bonding. During speech, mechanosensory neurons in the tongue and hard palate provide sensory inputs needed to produce vowels and sibilants (Hickok, 2012; Niemi et al., 2006; Niemi et al., 2002). Furthermore, lingual somatosensory neurons are important for execution of parental bonding behaviors, such as maternal licking behavior in rodents (Stern & Johnson, 1989) Despite these essential functions, only a handful of studies have interrogated the neural and cellular substrates of somatosensation in the tongue.

Several studies have investigated mechanisms of temperature and chemosensation in the oral cavity, but few have addressed the cellular and molecular mechanisms of mechanosensation in lingual neurons (Dickman et al., 1987; Donnelly et al., 2018; Leijon et al., 2019; Poulos & Lende, 1970a, 1970b; Robinson, 1992; Smith & Robinson, 1995; Yarmolinsky et al., 2016). This is particularly important as the tip of the tongue is exquisitely mechanosensitive with acuity comparable to that of the fingertip (Miles et al., 2018; Van Boven & Johnson, 1994). In previous studies, we reported the neurochemistry and morphology of peripheral neurons innervating the tongue in mice and humans, including mechanosensory axons in fungiform and filiform papillae that express the mechanically gated ion channel Piezo2 (Moayedi et al., 2018; Moayedi et al., 2021). The anatomy of these mechanosensory terminals are distinct from Piezo2-positive afferents in skin and other tissues; therefore, their functions in sensation are unknown. Previous electrophysiological studies identified a variety of physiological types of neurons innervating the tongues humans, cats and mice (Biedenbach & Chan, 1971; Grayson et al., 2019; Robinson, 1992; Smith & Robinson, 1995; Trulsson & Essick, 1997, 2010). In humans and cats, the majority of tongue mechanoreceptors are rapidly adapting, a typical property of Meissner corpuscles, indicating that the tongue is tuned for the detection of moving stimuli (Biedenbach & Chan, 1971; Mountcastle et al., 1967; Robinson, 1992; Trulsson & Essick, 1997, 2010). In addition, mouse geniculate ganglion mechanoreceptors selectively respond to moving stimuli and not to pressure (Yokota & Bradley, 2017). The trigeminal ganglion is well known to encompass a rich diversity of chemo-, thermo- and mechanoreceptors ; however, few studies have specifically analyzed lingual trigeminal neurons in mice (Dvoryanchikov et al., 2017; Nguyen et al., 2019; Nguyen et al., 2017; Wu et al., 2018; Yokota & Bradley, 2017; Zhang et al., 2019). Thus, open questions remain about whether the tongue is innervated by distinct types of mechanoreceptors capable of encoding diverse features of texture, whether rare neuronal groups are present, and whether mechanosensory neurons are molecularly distinguishable.

Here we use massively parallel in vivo imaging to investigate the functional, molecular and anatomical properties of tongue-innervating trigeminal ganglion neurons. We developed a hierarchical clustering approach to classify temporal response patterns of mechanosensory neurons, which provides an unbiased way to perform calcium imaging analysis. Furthermore, we identified two rare groups of mechanoreceptors that have not been previously described in mice.

## Methods

### Animal assurance statement

All animal experiments were conducted in accordance with Columbia University’s Animal Care and Use Policies. Experiments were performed with male and female mice, 4-11 months old.

### Mouse lines

Mouse lines used in this study included *Wnt1*^*Cre*^ (*129S4*.*Cg-E2f1*^*Tg(Wnt1-cre)2Sor/J*^, Jackson labs RRID:IMSR_JAX:022137), *Ret*^*CreERT2*^ (*Ret*^*tm2(cre/ERT2)Ddg*^, MGI:4437245), *Ntrk3*^*CreERT2*^ (*Ntrk3*^*tm3*.*1(cre/ERT2)Ddg/J*,^ RRID:IMSR_JAX:030291), *Vglut3*^*Cre*^ (*Slc17a8*^*tm1*.*1(cre)Hze*^, RRID:IMSR_JAX:028534), *Rosa26*^*Ai95*^ (*B6J*.*Cg-Gt(ROSA)26Sor*^*tm95*.*1(CAG-GCaMP6f)Hze*^*/MwarJ*, RRID:IMSR_JAX:028865), *Rosa26*^*mT/mG*^ (*B6*.*129(Cg)-Gt(ROSA)26Sor*^*tm4(ACTB-tdTomato,-EGFP)Luo/J*^, RRID:IMSR_JAX:007676), *Tau*^*eGFP*^ (*B6*.*Cg-Mapt*^*tm1(EGFP)Klt Tg(MAPT)8cPdav/J*^, RRID:IMSR_JAX:005491).

### Tamoxifen injections

For tamoxifen inducible lines, 150 mg/kg tamoxifen was injected intraperitoneally once between P21-P30. Tamoxifen was dissolved in 10% ethanol diluted in corn oil.

### In vivo imaging

*In vivo* calcium imaging of tongue-innervating trigeminal neurons was conducted in anesthetized mice as previously described (Yarmolinsky et al., 2016). In brief, mice were deeply anesthetized with ketamine (100 mg/kg) and xylazine (10 mg/kg). Mice were head-fixed to a stable rod, tracheotomized, and hemispherectimized to expose the trigeminal ganglia. Once hemostasis was achieved, the tongue was affixed to a platform. A force-controlled indenter (300C Dual Mode Muscle Lever; Aurora Scientific, Aurora, Canada) with a 2 mm tip was used to apply mechanical stimuli to the tongue. Brushing of the tongue was used to identify the region with the dense tongue-stimulation evoked responses. Brushing was applied using a wire cell culture loop 10 times from posterior to anterior followed by 10 times from anterior to posterior. Cold and cool flow stimuli were applied to the tongue with a 1 ml handheld pipette. Images were collected at 10 Hz using a Scientifica MultiPhoton In Vivo SliceScope equipped with an Olympus Plan N 4x (NA 0.10) lens. Illumination was provided by a CoolLED pE-300 (Andover, UK). Imaging was performed using Ocular (Teledyne Photometrics; Tucson, AZ). Image capture, LED shuttering, and pressure applications were synchronized using a Digidata 1550B (Molecular Devices; San Jose, CA).

### In vivo imaging analysis

Motion correction was performed using NoRMCorre (Pnevmatikakis & Giovannucci, 2017).Neurons were picked for analysis using the Cell Magic Wand plugin in ImageJ. Image files were converted to ΔF using image subtraction, and neurons were picked as cells with calcium fluctuations during any stimulation window. Neurons were excluded from analysis if they did not show calcium peaks during a stimulation window, or if they were clearly out of focus. After neurons were assigned Regions of Interest (ROIs), neuropil extraction and ΔF/F calculation was performed using Matlab code modified from (Ghitani et al., 2017) with the median fluorescence for the entire trace used as F_0_. Custom Matlab code was used to filter traces, identify peaks, remove spontaneous firing neurons, and assign to response group (e.g. pressure, brush). Traces were lowpass filtered using a 4^th^ order polynomial, with passband frequency of 2, and passband ripple of 0.5. Filtered traces were then smoothed using a Savitzky-Golay filter function with a 1st order polynomial over a 9-sample span. Peaks were then identified using the find peaks function in Matlab with a peak threshold of 1.5 ΔF/F. Peaks were aligned to stimulation windows with buffering time at the end of stimulus application to account for off responses. Neurons were defined as spontaneously firing and removed from analysis if the number of peaks outside of the stimulation windows was greater than the number of peaks inside of the stimulation widows divided by half of the total number of peaks:

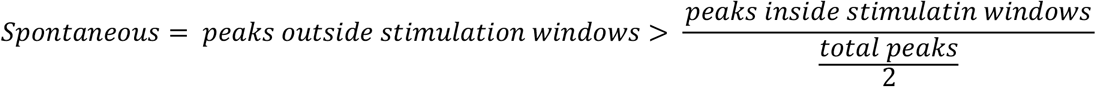

Cell areas were calculated using ROI assigned in ImageJ.

### Cell clustering

Clustering analysis was performed to identify mechanosensory neuron subtypes based on their responses to pressure stimuli using R. Collected time-series ΔF/F traces from the neuropil extraction program for all cells were first cleaned before clustering. More specifically, the high-frequency noise was removed using the Butterworth low-pass filter with a frequency of 0.05. The baseline drift was corrected by deriving trends for intervals between pressure stimuli using the Butterworth low-pass filter with frequency of 0.005 and connecting inter-stimulus interval trends using spline interpolation. Responses to pressure stimuli were detected if they were above a threshold of twice the highest value among the first 60-second unstimulated period. To offset differences in fluorescence due to imaging sessions or inherent difference in GCaMP expression, detected responses of all cells were normalized to share the same highest value. As time intervals between two pressure stimuli was 50 seconds long, 40 seconds of each inter-stimulus intervals was removed to condense signals to pressure responses only

The preprocessed, neuropil extracted ΔF/F traces were clustered in a two-layer structure. In the first layer, hierarchical clustering was conducted using the Ward1 clustering linkage criterion and the Euclidean distance of Fourier coefficients as the distance measurement (Mori et al., 2016; Murtagh & Legendre, 2014). The number of clusters was determined by incrementally increasing from a small number until no clusters with new patterns could be separated out. In the second layer, the same hierarchical clustering scheme was applied to each cluster obtained in the first layer to examine subclusters. In layer 2, de-noised and baseline corrected calcium traces without response detection and normalization were analyzed. Per first-layer cluster, if all subclusters in the second layer exhibited the same pattern, that first-layer cluster would be assigned as one individual neuron type. Otherwise, subclusters with new response patterns would be separated from the first-layer cluster and assigned as new cluster types. In addition to the above multi-layer hierarchical clustering, single-layer clustering and partitioning clustering were also tested for comparison.

### Peak analysis

Peak values were analyzed using custom-written code in MATLAB. In brief, mean grey value was extracted for each ROI to obtain raw fluorescence traces. Pressure and brush traces for each ROI were detrended by removing a 2nd or 3rd order polynomial trend (detrend() function in MATLAB). Traces were separated into pressure and brush stimulus windows (frames 1 to 3550 and frames 3551 to 1680, respectively) for detrending. Following this step, traces were separated into stimulus-response windows by extracting the fluorescence values (F) from the 5 seconds prior to and 20 seconds after stimulus onset (25 seconds total). The resulting traces were normalized to baseline (F_0_), defined as the average fluorescence from the 5 seconds prior to stimulus onset, and subsequently smoothed using a 3-point moving average filter. Max peak was defined as the maximal change in normalized fluorescence (NΔF/F) from baseline within the first three seconds following stimulus onset. Steady-state peak values were calculated by averaging the normalized fluorescence during the last two seconds of stimulation.

### Histology

Immunohistochemistry was performed as previously described (Moayedi et al., 2018). Tongues were flash frozen in OCT (Tissue-Tek) in liquid nitrogen. Trigeminal ganglia were fixed in 4% paraformaldehyde for 2 hours, washed in PBS, and submerged in 30% sucrose. Tissue cryosections (25 µm) were prepared on slides. Sections were dried at 37°C for one hour, and tongue sections post-fixed in 4% paraformaldehyde for 15 minutes. Slides were washed in PBS and incubated in NGS+PBST (PBS, 0.3% triton-X 100) with 5% normal goat serum (NGS) at room temperature for 1 hour. Sections were then incubated overnight in primary antibody mixed in NGS+PBST at 4°C. The next day, slides were washed three times in PBST and then incubated for two hours in secondary antibody mixed in NGS+PBST. Following this, slides were washed five times in PBST, and then mounted in Fluoromount-G with DAPI (Southern Biotech).

Antibodies used in this study were chicken anti-GFP (1:1000 Abcam, ab13970, lot GR236651-25, RRID:AB_300798), Rabbit anti-beta 3 tubulin (1:3000, Abcam, ab18207, lot GR3221401-3, RRID:AB_444319), rat anti-keratin8 (1:100, Developmental Studies Hybridoma Bank, supernatant, RRID:AB_531826), Alexa488 anti-chicken (1:1000, ThermoFisher, A-11039, AB_2534096), Alexa594 anti-rat(1:1000, Fisher Scientific, A11007, RRID:AB_141374), Alexa647 anti-rabbit(1:1000, Fisher Scientific, A21244, RRID:AB_2535812)

### Confocal Microscopy

Histology was imaged using a Nikon SP8 confocal microscope using LasX software. Images were taken with either a 10x NA=0.40 air lens or 40x oil immersion lens with NA=1.3. Images were taken at 2048×2048 pixels with 2x line averaging and Z-step size of 1 µm. Analysis was performed in ImageJ. Images were prepared for presentation in Adobe Photoshop by adjusting the threshold across the entire image.

### Quantification of Cre+ trigeminal neurons

Confocal images of trigeminal ganglia from each Cre line (N=2-3 image/ganglia, 6 ganglia from 3 mice per strain) were analyzed for GFP expression. Images from a single ganglion were collected from sections located at least 100 µm apart. For each genotype, images were acquired using the same confocal acquisition settings across all experiments. Neuron regions of interest were picked in imageJ based on β3-tubulin expression and presence of a discernable nuclei. At least 200 neurons were counted from each ganglion. A 2-pixel Gaussian blur and threshold was applied to include the top 3% of pixels. Regions of interest in which the majority of pixels were GFP+ were counted as Cre+ cells.

### Statistical analysis

GraphPad Prism 9 was used for graph visualization and statistical analysis. Data were tested for normality and then teste with either One-way ANOVA with Tukey’s post hoc or Kruskal Wallis test with Dunn’s multiple comparisons was used to compare means of multiple groups. Categorical data were analyzed with Chi square tests. Linear regressions were used to assess differences in force-response curves.

## Results

To analyze tongue-innervating trigeminal neurons, we adapted methods for *in vivo* calcium imaging of the trigeminal ganglia(Yarmolinsky et al., 2016). Previous studies have shown that trigeminal neurons have robust responses to cold stimulation of the oral cavity (Leijon et al., 2019; Yarmolinsky et al., 2016). Thus, we first tested whether tongue-innervating trigeminal neurons displayed cool flow and pressure evoked calcium responses using mice that express GCaMP6F in all trigeminal cells (*Wnt1*^*Cre*^, *Rosa26*^*Ai95*^). The tongue was gently pulled out of the mouth using blunt forceps and stabilized to a platform, allowing stimulation of the tongue independent of the rest of the oral cavity (**Figure 1A**). We applied five discrete pressure stimulations using a force-controlled indenter (range 0.09-0.44 N, or 29–140 kPa), followed by room temperature (RT) and cold flowing water (**Figure 1B**). Individual neurons responded to flowing liquid, pressure, or were polymodal with both pressure and flow-evoked responses (**Figure 1C-D**). Pooled data from hundreds of neurons show that distinct subsets of neurons respond to RT or cold flowing water, pressure, and both stimulus modalities (**Figure 1E**). The majority of responding neurons were RT/cold flow sensitive (83%, **Figure 1F**). These neurons tended to respond to both RT and cold flow stimuli, with greater response magnitude for cold stimulation compared to RT, consistent with previous studies of oral thermoreceptors (Leijon et al., 2019; Yarmolinsky et al., 2016). Around 17% of trigeminal tongue-innervating neurons that responded to stimulation in this paradigm were pressure sensitive (**Figure 1E-F**). Of these, less than half responded only to pressure, while the rest were polymodal, responding both to pressure and RT/cold flow, similar to previous findings in cats (Biedenbach & Chan, 1971; Robinson, 1992). We also compared the cell body sizes of response groups, and found that tongue-innervating trigeminal neurons that were responsive only to pressure were significantly larger than both the RT/cold flow responsive neurons and multimodal neurons (**Figure 1G**). Collectively, these data demonstrate that tongue-innervating trigeminal neurons include thermosensory, mechanosensory, and polymodal neuron classes.

**Figure 1.**
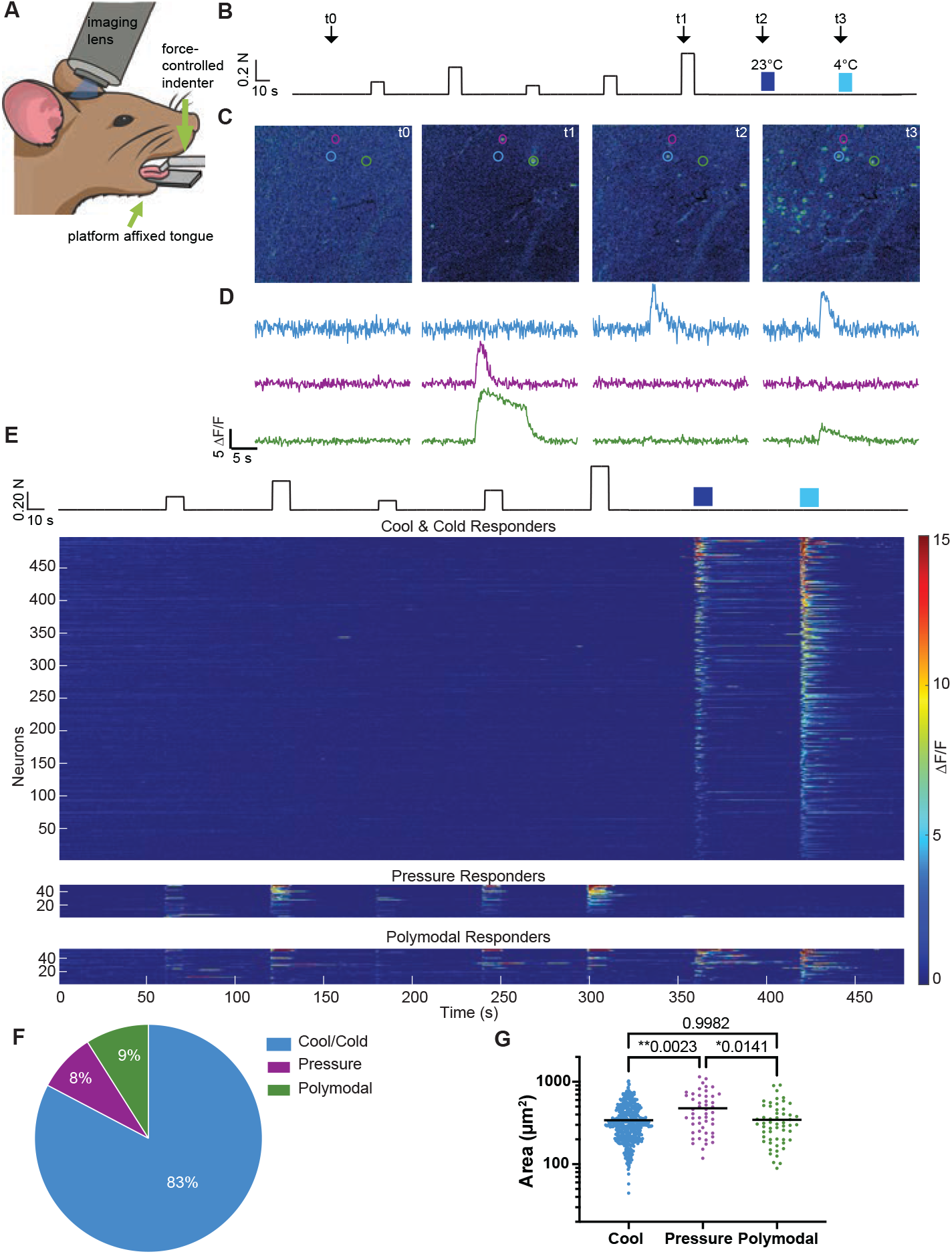
Somatosensory responses in tongue-innervating trigeminal neurons. **A**. Mice were head-fixed, craniotomized, and brain aspirated to reveal trigeminal ganglia. The tongue was then affixed to a platform and gently pulled outward to mechanically isolate it from the rest of the oral cavity. A force-controlled indenter was used to stimulate the tongue on the ipsilateral side as the exposed trigeminal ganglion and liquid stimuli were applied by hand to the tongue. **B**. The stimulation paradigm is shown. Force controlled indention was applied to the tongue between 0.09 N-0.44 N. After 60 s of baseline collections, stimulations were applied for 10 s, followed by 50 s rest before the next stimulation. This was followed by application of room temperature (RT) water (23°C) followed by cold water (4°C). Times for sample images are indicated by arrows and labeled as t0-t3. **C**. Sample images from an experiment are shown at baseline (t0), 0.40 N indention (t1), RT water (t2) and cold water (t3). Cells are circled that respond to cool/cold flow only (blue), pressure only (purple), and pressure and cold (green). **D**. Sample traces from cells circled in C. **E**. Summary data including responses from 601 neurons (N=17 *Wnt1*^*Cre*^, *Rosa26*^*Ai95*^ mice). Neurons were clearly separated into groups that responded to cool and/or cold only, pressure only, or both (polymodal). **F**. Distribution of neurons in each response category. **G**. Soma size was calculated for each neuron response class. Neurons that respond to pressure only were significantly larger than cool/cold or multimodal responders (Welch’s ANOVA P=0.0029 with Dunnett’s multiple comparisons, grey bar denotes median).

Most oral behaviors (e.g., feeding, vocalization, grooming) involve the movement of the tongue causing fluctuations in force across the mucosal surface; therefore, we posited that the rodent tongue might be equipped with mechanoreceptors tuned to detect moving stimuli. To test this hypothesis, we tested whether tongue mechanoreceptors classes preferentially responded to sustained pressure or to dynamic brush stimulation (**Figure 2**). Almost all neurons were brush sensitive (95%). Of these, 72% responded only to brush and 23% responded both to brush and pressure. The remaining 5% of neurons were sensitive to pressure only (**Figure 2B-E;** N=313 neurons from 13 mice). These data are in excellent agreement with previous electrophysiological studies from cat lingual afferents and mouse cutaneous afferents (Bai et al., 2015; Biedenbach & Chan, 1971; Robinson, 1992). Among pressure-responsive neurons, we noticed that neurons responded with subjectively different decay kinetics; neurons that responded only to pressure tended to do so with sustained calcium increases throughout the stimulus duration, whereas brush and pressure-responsive neurons tended to show transient increases in fluorescence. Moreover, brush-only responding neurons also had significantly smaller somata than pressure-only responding neurons (**Figure 2F**). Thus, as in the skin, the tongue is innervated by submodalities of mechanoreceptors with distinct temporal response patterns to sustained pressure, sensitivity to different modalities of touch, and somatal diameters.

**Figure 2.**
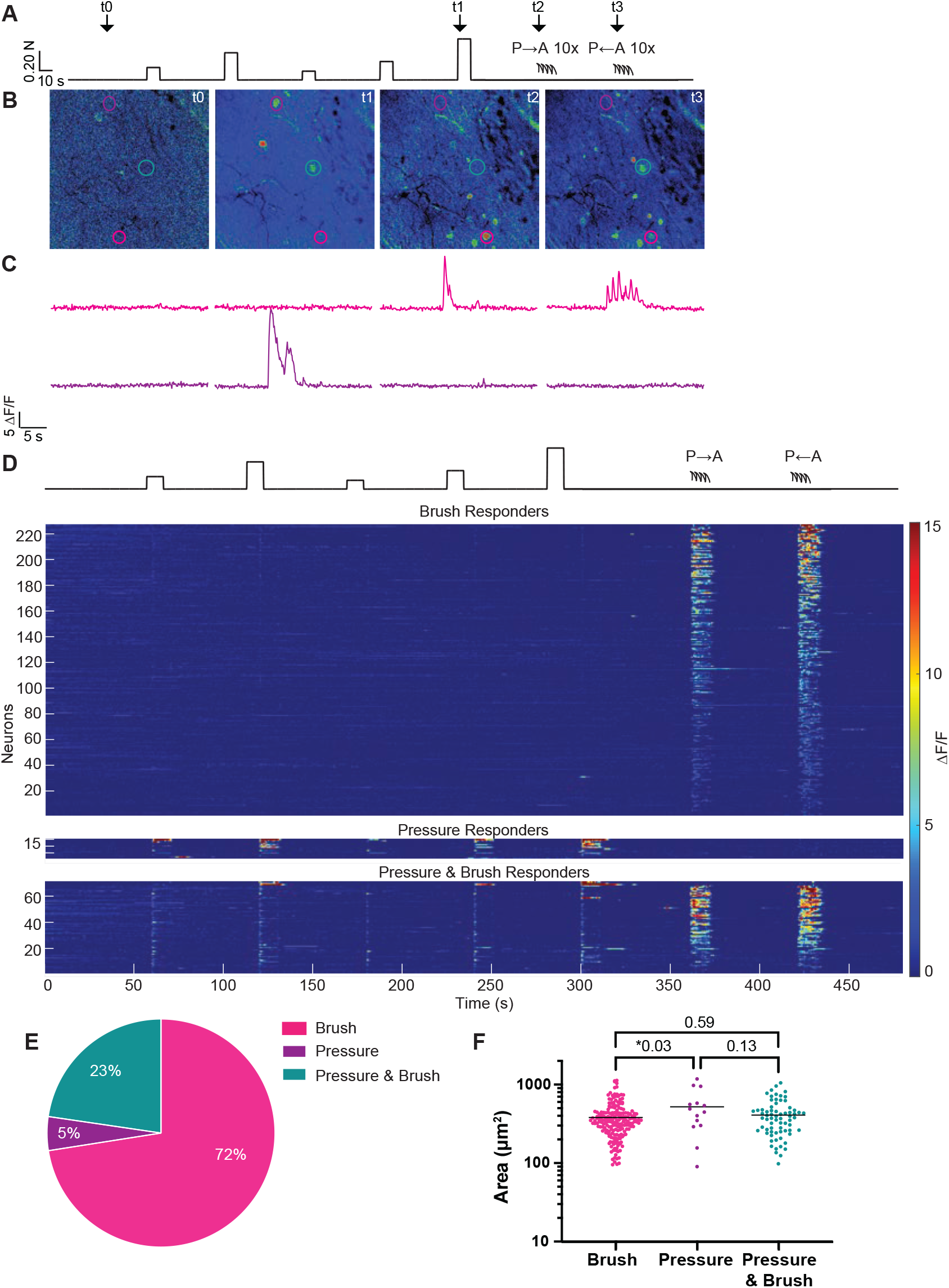
Tongue-innervating trigeminal mechanoreceptors are brush and/or pressure responsive. **A**. Stimulus paradigm is shown. Force controlled indention was applied as in **Figure 1** followed by 10 tongue brushes from posterior (P) to anterior (A) then A to P. **B**. Sample images from a single experiment are shown at baseline (t0), 0.4 N indention (t1), P to A brush (t2) and A to P brush (t3). Cells are circled that respond to brush only (pink), pressure only (purple), and pressure and brush (turquoise). **C**. Sample traces from cells circled in B. **D**. Summary data from 313 responding neurons from 13 mice **F**. 73% of responding neurons responded only to brushing, 5% only responded to pressure, and 23% were responsive to both. **G**. Pressure responsive neuron somas were significantly larger than brush-only responsive neurons. (one-way ANOVA P=0.0346 with Tukey’s post-hoc, grey bar denotes median).

In the skin, genetic markers label subsets of mechanoreceptors with different functional properties; therefore, we screened transgenic Cre mouse lines that label cutaneous low-threshold mechanoreceptors to determine whether they also mark tongue-innervating trigeminal mechanoreceptors. Cre-driver lines were crossed with mouse reporter lines that express membrane-bound GFP (Hippenmeyer et al., 2005; Muzumdar et al., 2007). We found that trigeminal ganglion neurons were labeled with three different Cre markers (**Figure 3A)**: the GDNF receptor Ret *(Ret*^*CreERT2*^ tamoxifen-induced at P21–30), vesicular glutamate transporter 3 (*Vglut3*^*Cre*^) and the NGF receptor *Ntrk3*^*CreERT2*^ (tamoxifen-induced at P21–30; (Bai et al., 2015; Griffith et al., 2019; Grimes et al., 2011; Lou et al., 2013; Luo et al., 2009). We quantified expression of each marker in the trigeminal ganglion and found that *Ret*^*CreERT2*^ labeled 31% of trigeminal neurons, *Vglut3*^*Cre*^ labeled 29% of trigeminal neurons, and *Ntrk3*^*CreERT2*^ labeled 10% of trigeminal neurons. Cre lines were then used to drive GCaMP6f expression in subsets of trigeminal neurons, and tested for their ability to visualize tongue-innervating mechanosensory responses. Ret+ neurons encompassed brush-, pressure-, and brush/pressure-sensitive subsets of mechanosensory neurons (**Figure 3B**). In response to sustained pressure, distinct temporal response patterns were also observed, with most neurons showing transient responses and some showing sustained responses. Vglut3-lineage neurons were also pressure/brush sensitive, but they lacked sustained responses to pressure. Ntrk3-expressing neurons were also sensitive to both pressure and brushing, and that pressure responses were a mixture of both sustained and transient types. By contrast, *Ntrk2*^*CreERT2*^, which marks a subset of rapidly adapting dorsal root ganglion neurons, and *Parv*^*Cre*^, which labels proprioceptors in dorsal root ganglion, showed no responses in trigeminal ganglia when the tongue was mechanically stimulated (Hippenmeyer et al., 2005; Rutlin et al., 2014). These data are consistent with previous reports that very few TrkB-positive trigeminal neurons innervate the tongue, and that the cell bodies of proprioceptors that innervate oral regions reside in the mesencephalic nucleus (Huang et al., 2021; Lazarov, 2007; Wu et al., 2018). Collectively, our results show that lingual mechanosensory neurons preferentially encode dynamic mechanical stimuli in mice. Although distinct temporal response patterns were observed, they did not correlate cleanly with the transgenic markers tested.

**Figure 3.**
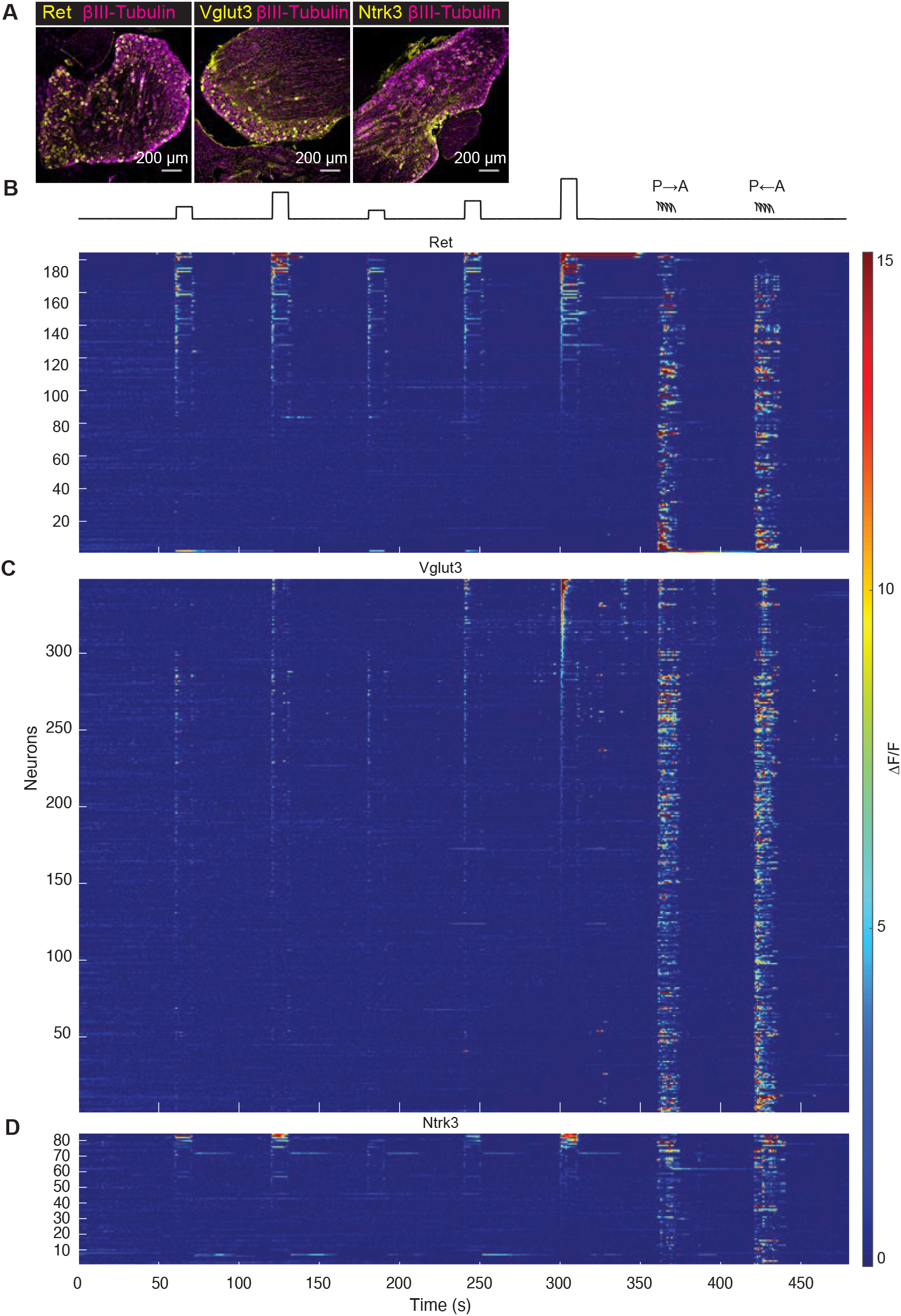
Tongue-innervating trigeminal mechanosensory neuron subgroups are distinguished by cre lines. **A**. Trigeminal ganglia of *Ret*^*CreERT2/+*^, *Rosa26*^*GCaMP6f/+*^; *Vglut3*^*Cre/+*^, *Rosa26*^*GCaMP6f/+*^; *Ntrk3*^*CreERT2/+*^, *Rosa26*^*GCaMP6f/+*^ animals reveal neuronal types marked by each line. **B**. Ret+ tongue-innervating trigeminal mechanosensory neurons are both brush and pressure sensitive. Distinct response dynamics of pressure sensitive neurons are apparent on heatmaps (184 neurons total from 7 mice). **C**. Vglut3 lineage tongue innervating trigeminal mechanosensory neurons are both brush and pressure sensitive. Pressure sensitive neurons primarily show transient responses to pressure (369 neurons from 10 mice). **D**. Ntrk3+ mechanosensory tongue innervating trigeminal neurons are both pressure and brush responsive. Pressure responsive neurons show a variety of response dynamics apparent by heatmap (90 neurons from 7 mice)

We next sought an objective method to determine how many distinct functional groups of mechanosensory neurons innervate the tongue. Somatosensory neurons are typically grouped based on their responses to discrete stimulus modalities (e.g., thermal, mechanical, noxious) or through subjective categorization of response properties (e.g., transient, sustained). These subjective grouping schemes are prone to experimenter bias and can fail when neurons exhibit intermediate response properties, or when rare populations exist. Thus, we sought to develop a scheme for unbiased clustering of mechanoreceptors to pressure. Data were combined from all mechanosensory neurons in *Ret*^*CreERT2*^, *Vglut3*^*Cre*^, and *Ntrk3*^*CreERT2*^ lines and blinded to mitigate experimenter bias. Combined time-series data were preprocessed before clustering, which included noise removal, baseline drift correction, peak detection, normalization to peak, and inter-stimulus interval removal (**Figure 4A**). We tested both partitioning and hierarchical clustering methods. Results from partitioning clustering were not stable across different runs. Clustering in this scheme also failed to distinguish subtle differences observed in traces with low-amplitude responses, as described below. By contrast, hierarchical clustering results were constant across repeated runs. Among multiple linkage criteria tested, Ward1 criterion resulted in the clear separation of unique clusters. We then applied a second iteration of clustering to each primary cluster to examine if additional groups were present. In this round, traces were only denoised and baseline corrected. Such less-processed data retrieved the information removed by response detection, which allows the identification of neurons that display reduced fluorescence in response to force. (**Figure 4A**). Initial hierarchical clustering identified four clusters with distinct temporal patterns (**Figure 4B**). When *Cl1, Cl2* and *Cl3* were individually analyzed in a second clustering round, response patterns of subclusters were indistinguishable within each cluster. By contrast, *Cl4* separated into four subclusters. Surprisingly, force steps suppressed fluorescence in two of these subclusters. Thus, we separated *Cl4* into a group with little or no fluorescence change (*Cl4a*) and those that showed fluorescence reductions during force steps (*Cl4b*). Overall, our hierarchical clustering approach identified five functional groups of tongue-innervating mechanosensory neurons based on response kinetics and amplitude.

**Figure 4.**
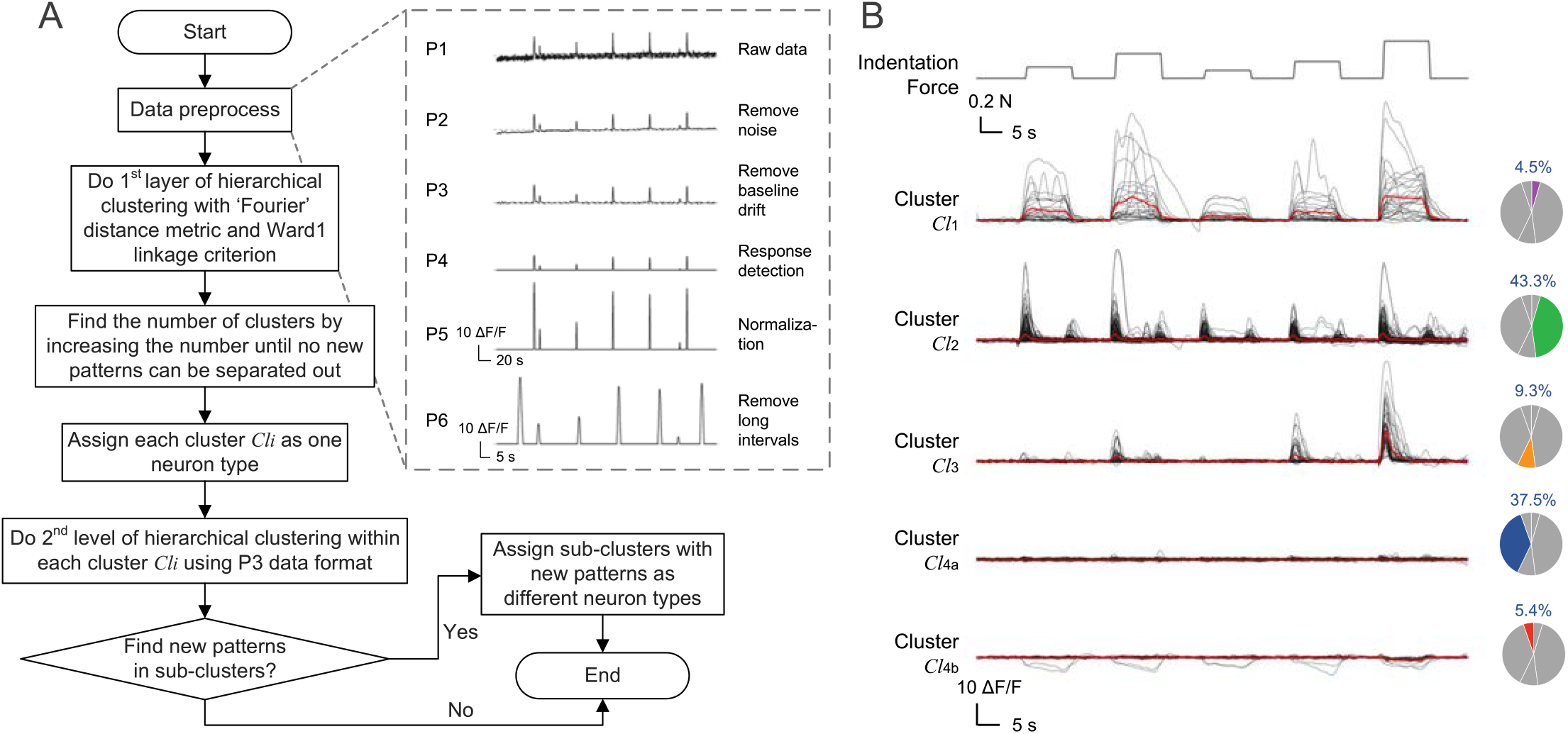
Hierarchical clustering approach to identify sub-populations of pressure-responsive neurons. **A**. Logic diagram depicting steps for hierarchical clustering. **B**. Hierarchical clustering revealed four distinct clusters of response types in the first layer. Pressure responses from each group are shown with average response in red. Within cluster 4, a significant number of neurons showed a negative going response, thus, these were separated into a distinct cluster.

We next quantitatively analyzed the response properties of neurons in each cluster (*Cl1-4b*, **Figure 5**). Clusters were unequally represented across the population, ranging from 4.5–43.3% of mechanosensory neurons (**Figure 5A**). Clusters displayed differences in average amplitudes and temporal kinetics to the highest force levels tested (**Figure 5B**) as well as median soma size (**Figure 5C**), indicating that biological differences within the population of tongue-innervating mechanosensory neurons were captured in the clustering approach.

**Figure 5.**
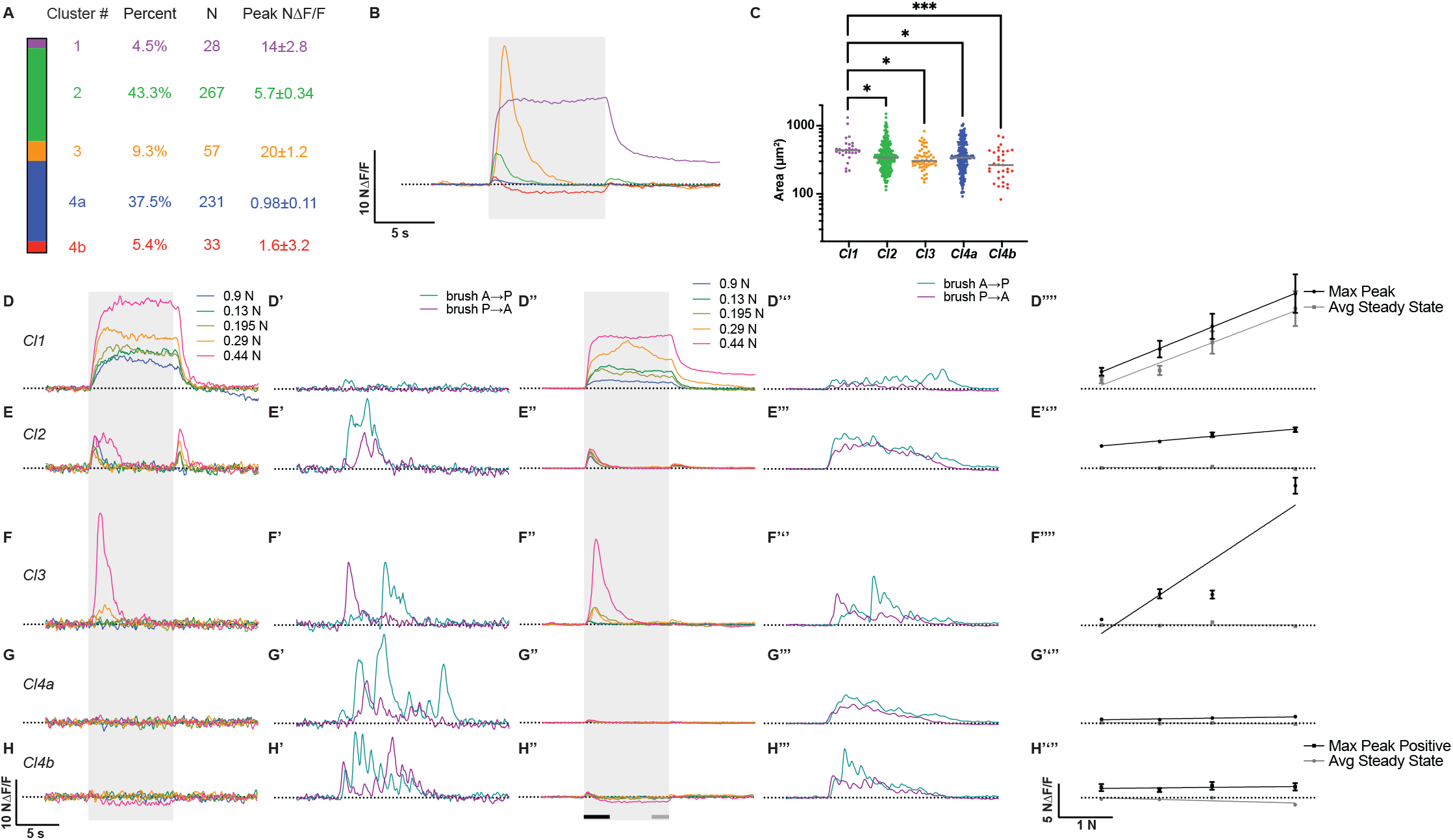
Clustering reveals five populations of tongue innervating trigeminal mechanoreceptors with distinct response dynamics and cell body sizes. **A**. Percent, number of neurons, and peak NΔF/F±SEM are shown for each cluster. **B**. Average responses (normalized change in fluorescence, NΔF/F) to maximum stimulation (0.44 N) for each cluster. Responses have subjectively distinguishable differences between magnitude and direction of response as well as kinetics of force response (shaded box indicates stimulation window). **C**. Cell areas for each cluster show that neuronal sizes are significantly different (Kruskal Wallis test P =0.0009, Dunn’s multiple comparisons *p<0.05, ***p<0.001). *Cl1* consists of the largest diameter neurons and *Cl4b* with the smallest (grey bar denotes median). **D-H**. Representative responses to each force stimulation for each cluster (shaded box indicates pressure stimulation window; **D**: *Cl1*, **E**: *Cl2*, **F**: *Cl3*, **G**: *Cl4a*, **H**: *Cl4b*). **D’-H’**. Representative responses to brushing from each cluster. **D’’-H’’**. Average responses to each force stimulation for each cluster. Bars below indicate stimulation windows from which maximum force responses (black) and average steady state responses (grey) are calculated. **D’’’-H’’’**. Average responses to brushing from each cluster. **D’’’’-H’’’’**. Maximum response (black) from the first 3 seconds and average steady state response (grey) from the last 2 seconds for each cluster are shown with linear fits to the population data. Dotted line denotes 0 NΔF/F. Mean±SEM. **D’’’’**. Cluster 1, maximum response to stimulation has a positive slope (slope=0.33 (NΔF/F)/N, R^2^=0.17, P<0.0001). Steady state responses to stimulation has similar positive slope (slope=0.31 (NΔF/F)/N, R^2^=0.19, P<0.0001). Maximum and steady state slopes are not significantly different (P=0.8491) **E’’’’**. Cluster 2, maximum response to stimulation shows a slight positive slope (slope=0.07 (NΔF/F)/N, R^2^=0.035, P<0.0001). Steady state responses to stimulation has a slight negative slope (slope=-0.0034 (NΔF/F)/N, R^2^=0.0055, P=0.015). Max and steady state peaks are significantly different (P<0.0001) **F’’’’**. Cluster 3, maximum response to stimulation shows a positive slope (slope=0.54 (NΔF/F)/N, R^2^=0.54, P<0.0001). Steady state responses to stimulation is not different from zero (slope=-0.0025 (NΔF/F)/N, R^2^=0.0034, P=0.0056). Max and steady state peaks are significantly different (P<0.0001) **G’’’’**. Cluster 4a, maximum response to stimulation shows a slightly positive slope (slope=0.012 (NΔF/F)/N, R^2^=0.015, P=0.0002). Steady state responses to stimulation has a slight negative slope (slope=-0.0046 (nΔF/F)/N, R^2^=0.043, P<0.0001). Max and steady state peaks are significantly different (P<0.0001). **H’’’’**. Cluster 4a, maximum positive response to stimulation has a flat slope (slope=0.008 (NΔF/F)/N, R^2^=0.0013, P=0.68). Steady state responses to stimulation has a slight negative slope (slope=-0.022 (NΔF/F)/N, R^2^=0.12, P<0.0001). Max and steady state peaks are not significantly different (P=0.14).

*Cl1* neurons (4.5%) showed large, sustained fluorescence increases during force steps, but weak responses to brush stimuli (**Figure 5D-D’’’**). We noted that these neurons had significantly larger somata that all other clusters (**Figure 5C)**. Analysis of force-response curves showed that the activity of *Cl1* neurons was positively correlated with force magnitude at both the initial peak and steady state (**Figure 5D’’’’**). Comparison of linear fits between peak and steady state responses show that the two slopes did not differ significantly. Thus, Cl1 neurons, likely slowly adapting touch receptors, encode force magnitude throughout the duration of 10-s pressure stimuli.

By contrast, *Cl2* neurons, which were most abundant in this study (43%), showed small, transient “on” responses at all force levels tested, and were robustly activated by brush (**Figure 5E-E’’’**). Some of these neurons also showed “off” responses, which were generally smaller than on responses. *Cl2* peak responses had a slightly positive slope to increasing forces, indicating that these neurons only weakly represent stimulus magnitude in their neuronal activity (**Figure 5E’’’’**). Steady state response amplitudes were near baseline. Together, these data indicate that *Cl2* neurons, like rapidly adapting mechanoreceptors, are tuned to report the presence, but not the magnitude, of dynamic stimuli.

*Cl3* neurons (9%) also showed transient responses to force steps; however, their stimulus-response relations were markedly different from other neuronal clusters. Peak fluorescence signals were similar among the low and intermediate forces tested, but these neurons exhibited large response amplitudes at 440 mN (**Figure 5F-F’’’**). Thus, this cluster showed a steep positive slope in their peak force-response relations (**Figure 5F’’’’**). Moreover, they responded more vigorously to force steps than brush. These response properties suggest that Cl3 neurons are high-threshold mechanoreceptors that respond preferentially to noxious mechanical stimuli.

Conversely, *Cl4a* neurons (38%) showed little or no calcium increases during force steps but responded vigorously to brush (**Figure 5G-G’’’**). Neurons in this cluster had a slightly positive initial force-response relation and were silent at steady state (**Figure 5G’’’’**). Thus, Cl4a neurons are brush receptors. Cl4b neurons (5%) showed similar brush sensitivity and very small fluorescence increases at the onset of force steps (**Figure 5H-H’’’)**. Unexpectedly, these neurons showed sustained fluorescence decreases at the highest force levels tested. Although the initial positive response showed a stimulus-response slope of zero, the magnitude of the steady state response was negatively correlated with force amplitude (**Figure 5H’’’’**). This surprising result indicates that a small population of tongue mechanoreceptors are progressively inhibited by increasing pressures but are robustly activated by brush.

We next compared the distribution of neurons derived from each Cre line in the full dataset with the distribution of genetic markers in each cluster to determine whether these molecular markers distinguish functionally defined neuronal classes (**Figure 6A**). The *Cl1* cluster, which responded best to force steps, comprised *Ntrk3*- or *Ret*-expressing neurons, with only 1/28 neurons of Vglut3 lineage. The distribution of molecular markers present in this cluster differed significantly from that of the population as a whole (Chi-Square test, p<0.0001). These data are consistent with expression of Ret and Ntrk3 in cutaneous slowly adapting mechanoreceptors (Bai et al., 2015; Funfschilling et al., 2004; Luo et al., 2009). *Cl2*, which showed transient responses at all forces, contained neurons from all three genetic marker strains at proportions similar to the parent population. *Cl3* neurons, which are putative mechanonociceptors, comprised 89% *Vglut3*-lineage neurons, 11% *Ret*-expressing neurons, and no *Ntrk3*-expressing neurons (Chi-Square test, p<0.0001). These data are consistent with known expression of Ret in high-threshold mechanoreceptors, although this is a novel finding for Vglut3-lineage neurons (Lou et al., 2013; Luo et al., 2009). *Cl4a* neurons included all three molecular markers, and were not significantly different from input. The *Cl4b* cluster contained almost entirely *Vglut3*-lineage or *Ntrk3*-expressing neurons, with only one Ret+ neuron. This distribution was particularly surprising because Ret expression is widely expressed in adult somatosensory neurons. Overall, the overlap in response properties between Cre-driver lines suggest that these genetic markers label overlapping subsets of trigeminal mechanosensory neurons.

**Figure 6.**
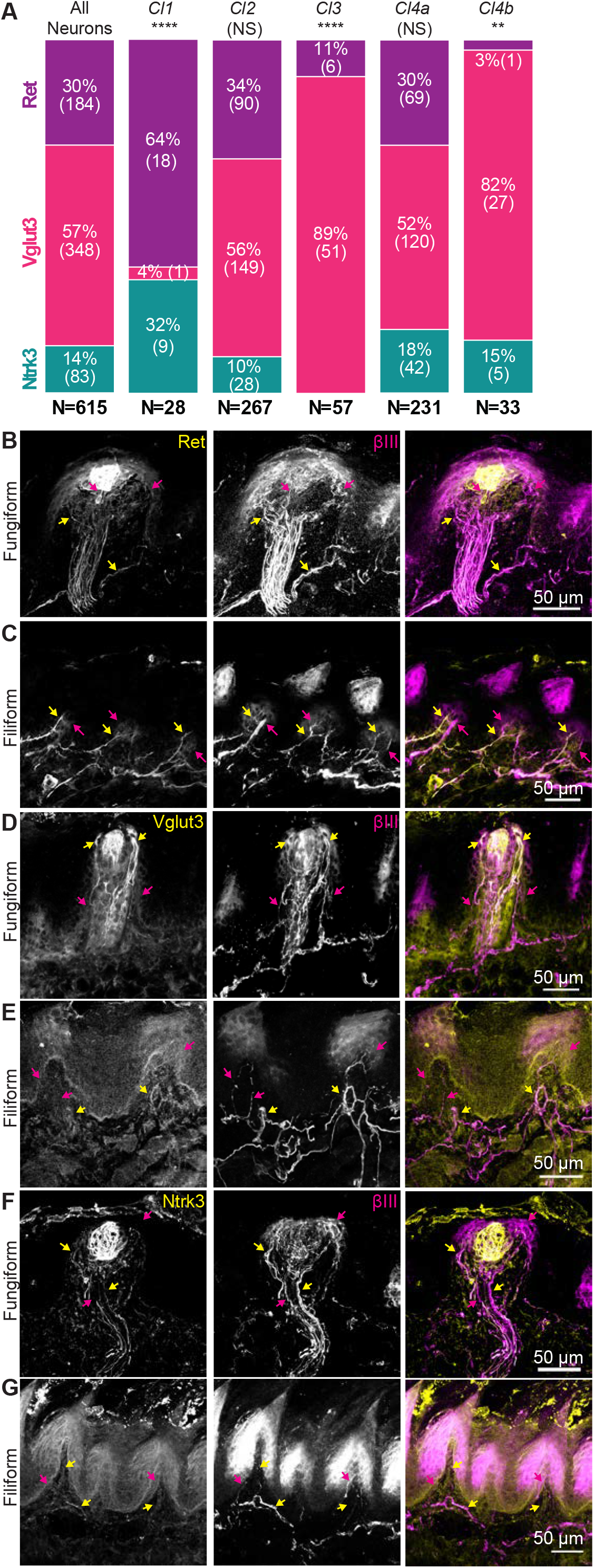
Molecular markers target clusters of tongue innervating trigeminal mechanosensory neurons. **A**. Distribution of neurons from each Cre line are shown in each cluster. Chi-square test were performed with the number of neurons analyzed from each molecular maker as the expected distribution (**p<0.01, ****p<0.0001). *Cl1, 3* and *4b* had significantly different representations of Cre lines compared to neurons analyzed. **B**. Afferents of *Ret*-expressing neurons commonly extend into fungiform papillae (yellow arrows), although some unlabeled neurons are also present (magenta arrows). **C**. *Ret*-expressing neuron afferents also commonly extend into fungiform papillae (yellow arrows), unlabeled afferents were also common (magenta arrows). ***D***. *Vglut3*-lineage neuron endings commonly extended into extragemmal epithelia in fungiform taste buds (yellow arrows). Non-*Vglut3*-lineage neurons were also commonly identified in fungiform papillae (magenta arrows). **E**. *Vglut3*-lineage afferents were occasionally found in filiform papillae (yellow arrows), although negative afferents were more common (magenta arrows). **F**. *Ntrk3* expressing neuron endings were identified as innervating fungiform papillae (yellow arrows). Negative endings were also present (magenta arrows). **G**. *Ntrk3* expressing afferents were commonly identified in filiform papillae (yellow arrows).

These results led us to ask whether molecular markers label distinct peripheral end organs in the tongue, and whether this could help us to assign end organs to neuronal clusters. We analyzed anatomical endings labeled by each Cre line using Cre-dependent membranous GFP expression (Hippenmeyer et al., 2005; Muzumdar et al., 2007). We found that end organs in both the fungiform and filiform papillae are labeled in all three mouse lines. (**Figure 6B-G**). Ret-expressing neurons were commonly found in neurons innervating the fungiform papillae, with some Ret-negative neurons also present (**Figure 6B**). Around half of filiform papillae were innervated by Ret+ neurons (**Figure 6C**). Vglut3-lineage neurons were commonly observed around fungiform papillae, often extending into the extragemmal epithelia (**Figure 6D**) and were occasionally identified in filiform papillae (**Figure 6E**). Ntrk3 expressing neurons extended afferents commonly in fungiform papillae in extragemmal space (**Figure 6F**), and in filiform papillae cores (**Figure 6G**). Collectively, we found no clear distinction between genetic markers among different end-organ morphologies.

## Discussion

Although tongue-innervating sensory neurons play a critical role in flavor, feeding, and social behaviors, the diversity of mechanosensory neurons populations innervating the tongue has long been uncharacterized. This is in part due to the technical limitations that arise when recording from lingual trigeminal neurons. Using *in vivo* trigeminal calcium imaging methods and a novel response-clustering paradigm, we identified five clusters of mechanosensory neurons with distinct response properties that innervate the mouse tongue. These neuron clusters differ in prevalence, cell body size, force-response relationship, and decay kinetics to force. Despite these differences in response profiles, distinct end organ structures correlating to response type are not clearly indefinable in these lingual afferents, suggesting that trigeminal tongue-innervating mechanoreceptors achieve similar response profiles to cutaneous afferents with a divergent end organ morphology.

We identified clusters of trigeminal tongue-innervating mechanosensory neuron types using a novel unbiased hierarchical clustering approach. Clustering response dynamics to pressure using a multilayer hierarchical approach offered significant advantages over the manual classification typically used in calcium imaging analysis including speed, objectivity of results, and identification of rare groups. In all, we identified five distinct clusters of tongue-innervating mechanosensory neurons based on their response profiles to ramp and hold pressure with response types reminiscent of known skin mechanoreceptors. *Cl1* included sustained responding neurons that encoded pressure well across the full range of stimuli given, similar to slowly adapting type responses of Merkel cell-neurite complexes and possibly Ruffini endings (Iggo & Muir, 1969; Johnson et al., 2000). Interestingly, this cluster was only weakly brush sensitive. These were both the largest and the rarest of the clusters identified. *Cl2*, the largest cluster, included neurons that had a transient response to stimulation active at stimulation-onset and, often, stimulation-offset for pressure and were also very responsive to brushing. These response characteristics greatly resemble the rapidly adapting (type of) response that is common for Meissner’s and Pacinian corpuscles (Mountcastle et al., 1967). *Cl3* neurons also displayed transient responses to stimulation, but were more active in high force ranges, resembling high-threshold mechanoreceptors (Bautista & Lumpkin, 2011). *Cl4a* neurons were primarily brush-sensitive, similar to a class of stroke-sensitive neurons identified in mouse hairy skin (Bai et al., 2015). The final cluster, *Cl4b*, displayed transient increase in activity due to pressure followed by a sustained force-encoding inhibition. This response type could either signify a spontaneously active neuron that has a transient increase in activity at high forces, followed by inhibition, or a neuron type that depletes internal calcium stores after excitation. *Cl4b* was also highly brush-sensitive and had the smallest soma size of all clusters. Collectively, these data suggest that trigeminal mechanosensory tongue-innervating neurons are capable of transducing a broad range of mechanical stimuli including innocuous, noxious, constant pressure, and movement. This is in stark contrast to what is known of geniculate mechanoreceptors, which have in previous studies been shown to respond only to moving stimuli and are genetically homogenous (Dvoryanchikov et al., 2017; Yokota & Bradley, 2017; Zhang et al., 2019).

Mechanosensory tongue-innervating trigeminal neuron clusters identified in this study are comparable to those identified previously in humans and cats (Biedenbach & Chan, 1971; Robinson, 1992; Trulsson & Essick, 1997, 2010). In humans, four low-threshold mechanosensory neuron groups have been identified using microneurography of the lingual nerve (Trulsson & Essick, 1997, 2010). These included rapidly adapting units, slowly-adapting units with regular firing, slowly adapting units with irregular firing, and deep mechanoreceptors with proprioceptive capabilities. Similar populations were identified in cats (Biedenbach & Chan, 1971; Robinson, 1992). Slowly adapting mechanoreceptors were the smallest response group identified in humans and cats; similarly, our study found that neurons with sustained responses to pressure (*Cl1*) were the least abundant class. Our study did not identify two clusters of sustained responding neurons, but this possibility is not excluded as calcium imaging is likely not able to distinguish the regularity of firing rate in the sustained phase. The majority of the neurons identified in humans and cats were rapidly adapting (Biedenbach & Chan, 1971; Trulsson & Essick, 1997, 2010), in agreement with our findings that *Cl2* was the most abundant cluster. In cats, an additional population of neurons was identified that was responsive to brushing but not pressure, similar to *Cl4b* in our study. This work did not definitively identify putative proprioceptors, as were identified in humans (Trulsson & Essick, 1997), but this possibility was not excluded by this experimental paradigm as classical proprioceptors are not known to have cell bodies in the trigeminal ganglion. This study identified two clusters of which correlates were not identified in humans or cats, *Cl3* and *Cl4b*. In prior work in humans, the force stimulation range was subthreshold to nociceptors, thus the methods used may not have been amenable to identifying high threshold mechanoreceptors (Trulsson & Essick, 1997, 2010). Furthermore, caveats of the methods used in human studies cited by the authors include a slight bias towards large-diameter neurons and spontaneously active neurons, collectively leading to potential under sampling of high threshold mechanoreceptors. In keeping with this, *Cl3* and *Cl4b* were the smallest diameter of all clusters identified in this study. In addition, due to the relatively small sample size that is feasible to achieve with microneurography, these rare units may not have been captured. One previous study recorded pressure-response dynamics in tongue-innervating mechanosensory neurons using mouse tongue-nerve preparations (Grayson et al., 2019). This study found that high-threshold mechanoreceptors were the most abundant class present in the tongue. The difference in findings between the two studies could be due to sensitization resulting from cell injection in the tongue, differences in neuronal cadres in athymic nude mice compared to immunocompetent animals, or due to biases inherent in methodologies used. These methodological differences and differences in presentation of the data make it difficult to directly compare our findings with this prior study. Together, we conclude that mice are equipped with a similar complement of low-threshold trigeminal tongue-innervating mechanosensory neuron classes as humans and cats, and have additional high threshold mechanoreceptor categories that were not previously described.

A question remains of how these clusters of tongue-innervating trigeminal neurons contribute to oral sensation. While behavioral analysis probing how these clusters contribute to mechanosensation have yet to be done, we can make inferences on their presumptive roles of clusters based on their activation range and kinetics of responses to force application. *Cl1* neurons are the only group that respond to force with a sustained increase in activity. This group was also active across the range of forces given, implying that it has a wide range of force coding abilities. This activity profile suggests that these would be good at detecting sustained forces, such as those felt when the tongue is resting against the roof of the mouth. *Cl2* neurons are active at low thresholds and only transiently respond to sustained force at onset and offset of stimulation. This response type is similar to the firing pattern of Meissner and Pacinian corpuscles, suggesting that these neurons would encode vibration and movement, such as those felt when the tongue is actively involved in feeding or speech. *Cl3* has a steep force-response relationship, and responds with a burst of activity at the onset of high pressure. These neurons are most likely a form of high threshold mechanoreceptor, and thus would encode painful pressure. *Cl4a* is primarily brush sensitive, so would likely only encode movement, like liquid flowing across the tongue, or what the mouse feels when it is grooming. Finally, *Cl4b* has a brief positive response to pressure followed by sustained reduction in activity and were highly brush-sensitive. How these neurons shape perception of force remains a mystery, but future functional studies can elucidate this through behavioral and electrophysiological analysis. Collectively, we found that the majority of neurons had transient responses to force and were brush sensitive, implying that the mouse tongue is best at encoding moving stimuli. This finding is consistent with those in humans and cats, and is in line with a role for the tongue in active sensing during dynamic tasks like feeding, speaking, and grooming (Biedenbach & Chan, 1971; Robinson, 1992; Trulsson & Essick, 1997). Previous studies found that the hard palate in mice is densely innervated with Merkel cells, indicating that this surface likely compensates for the tongue’s scarcity in pressure encoding, sustained responding neuronal types (Moayedi et al., 2018; Nunzi et al., 2004).

Mapping of molecular markers to lingual mechanosensory neuron groups reveals insights into how these functional classes compare to skin-innervating mechanoreceptors. Ret is expressed in around 60% of dorsal root ganglion neurons including non-peptidergic nociceptors, Meissner corpuscles, Pacinian corpuscles, and lanceolate endings (Luo et al., 2009). In oral tissues, Ret has been shown to be expressed in tooth pulp afferents (Donnelly et al., 2019) and in geniculate ganglion neurons innervating fungiform papillae (Donnelly et al., 2018). Only one cluster did not include Ret+ cells, *Cl4b*, thus molecular identification of this cluster can be performed through transcriptomic analysis of Ret-mechanoreceptors in the trigeminal ganglion in future studies. Vglut3 has previously been shown to be expressed transiently in Merkel cell afferents and persistently in C-LTMRs and a second class of unmyelinated free nerve endings innervating the skin (Lou et al., 2013). In lingual mechanoreceptors, Vglut3-lineage neurons were predominantly included in Cl2-4b. These include neurons with properties of transient LTMRs (*Cl2*), high threshold mechanoreceptors (*Cl3*), brush responsive neurons (*Cl4a*), and neurons with a sustained negative response after high force stimulation (*Cl4b*). These data suggest that the tongue may have further diversification of mechanosensory Vglut3-lineage neuron types compared to skin-innervating mechanoreceptors. Ntrk3 is expressed in Merkel cell afferents, A-beta field LTMRs, and a subset of free nerve endings (Bai et al., 2015). In adults, around half of adult TrkC+ DRG neurons are Ret+. Accordingly, we found that neurons from *Ntrk3*-expressing neurons were present in all lingual clusters except *Cl3*, the high threshold mechanoreceptor group. Collectively, this strategy identified 3 clusters that were distinguishable by the absence of a molecular marker. In future studies, transcriptomic and intersectional genetic approaches can be used to definitively identify and manipulate these neuronal clusters to identify their functions in lingual mechanosensation.

In the tongue, peripheral mechanosensory end organs do not have morphologies that are consistent with those of known mechanoreceptors of the skin, making inferences of how endings relate to function speculative. We sought out to identify whether specific end organs correlated with any of the genetic markers that we investigated in this work. We found that neuronal afferents from all three molecular markers were present in both the filiform and fungiform papillae without any clear end organ structures to delineate afferent types. Further complicating this quest to identify end organs associated with the populations identified in this study is that fungiform papillae are also innervated by neurons with cell bodies in the geniculate ganglia, some of which express Ret and TrkC (Donnelly et al., 2018; Matsumoto et al., 2001). the best of our knowledge, expression of Vglut3-lineage neurons has not been investigated in the geniculate ganglion. Thus, future studies will need to disentangle whether these molecularly labeled mechanosensory end organs in the fungiform papillae are derived from trigeminal or geniculate ganglia, and whether these are similar populations of mechanoreceptors in each nucleus.

## Acknowledgments

We thank David Ginty and Meenakshi Rao for providing *Ret*^*CreERT2*^ mice, David Yarmolinsky for training on trigeminal imaging procedures, and Alexander Chesler and Marcin Szczot for providing neuropil extraction code. We thank Dr. Rachel Clary and members of the SENse lab at UC Berkeley for useful discussions on this project. This work was supported by Berrie Foundation Initiative on the Neurobiology of Obesity (EAL, YM); Thompson Family Foundation Initiative on Chemotherapy Induced Peripheral Neuropathy and Sensory Neuroscience (YM); and NIH NINDS R01NS105241 (GJG, EAL), NIH NIDCD R21DC018898 (YM).

## Notes

**Conflict of interest statement.** The authors have no conflict of interests to report.

### Competing Interest Statement

The authors have declared no competing interest.

